# Cytokine Dynamics and Oxidative Stress in Host Cells Stimulated with Drug-Resistant and Sensitive *Mycobacterium tuberculosis* Isolates

**DOI:** 10.1101/2024.12.02.626310

**Authors:** Kavitha Kumar, Prashant Giribhattanavar, B. K. Chandrasekhar Sagar, Shripad A. Patil

## Abstract

**Background:** The immune response to *Mycobacterium tuberculosis* (M. tuberculosis) is central to the pathogenesis of tuberculosis (TB), yet the immune dynamics induced by drug-resistant strains remain underexplored. Understanding the host’s immune response to both drug-sensitive and drug-resistant *M. tuberculosis* isolates is crucial for elucidating the mechanisms of pathogenesis and resistance. This study aims to assess the cellular immune responses, including PBMC proliferation, cytokine secretion (IL-4 and IL-17a), and reactive oxygen species (ROS) production in response to live drug-sensitive and drug-resistant *M. tuberculosis* clinical isolates.

**Methods:** Peripheral blood mononuclear cells (PBMCs) from PPD-negative and PPD-positive healthy volunteers were stimulated with live *M. tuberculosis* isolates, including MDR, SI-resistant, and sensitive strains. The immune responses were assessed by evaluating cell proliferation, secretion of IL-4 and IL-17a cytokines, and ROS production over a 9-day period.

**Results:** PBMCs from PPD-positive individuals exhibited a higher proliferative response compared to PPD-negative individuals, indicating more robust immune memory. IL-4 secretion was low but varied among samples, with higher levels observed in response to MDR isolates, suggesting a potential role in immunopathology. IL-17a levels increased over time, particularly in PPD-positive individuals, and MDR strains elicited a stronger response than sensitive isolates. ROS production was significantly elevated in response to resistant strains, reflecting the host’s oxidative defense mechanisms.

**Conclusion:** This study demonstrates distinct immune responses to drug-resistant *M. tuberculosis* isolates, with variations in cell proliferation, cytokine secretion, and ROS production. These findings provide insights into the immune dynamics during infection with resistant strains and underscore the importance of genotype-environment interactions in TB pathogenesis.

**Graphical Abstract:** 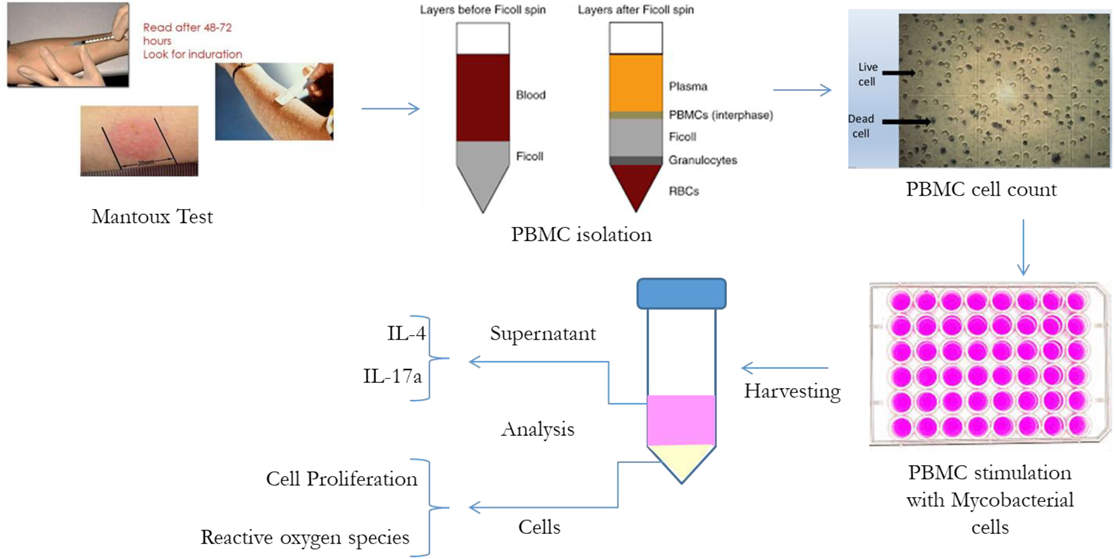

## Introduction

Tuberculosis (TB), caused by *Mycobacterium tuberculosis*, remains one of the leading causes of death globally, despite significant advances in diagnosis and treatment (Alsayed et al., 2023). In 2022, the World Health Organization (WHO) reported an estimated 10.6 million new TB cases and 1.6 million TB-related deaths globally, with an increasing burden of drug-resistant strains complicating management (Bagcchi 2023). Drug-resistant TB, particularly multidrug-resistant (MDR) and extensively drug-resistant (XDR) *M. tuberculosis*, poses a major threat to public health, requiring prolonged and often less effective treatment regimens (Lange et al., 2014). The emergence of these strains underscores the urgent need for a more comprehensive understanding of the host-pathogen interactions that drive disease progression and treatment failure (Sachan et al., 2023).

The immune response to *M. tuberculosis* is a complex process that involves both innate and adaptive immune mechanisms (de Martino et al., 2019). Upon infection, the pathogen is initially encountered by innate immune cells, such as macrophages and dendritic cells, which attempt to engulf and eliminate the bacilli (Lerner et al., 2015). These cells trigger the production of reactive oxygen species (ROS) and pro-inflammatory cytokines as part of the defense mechanism. ROS production is a key feature of the oxidative burst, wherein phagocytes, such as macrophages, produce superoxide radicals, hydrogen peroxide, and hydroxyl radicals to kill the bacteria (Juan-García et al., 2017). However, *M. tuberculosis* has evolved several mechanisms to resist oxidative stress, including the production of antioxidant enzymes like catalase-peroxidase (KatG) and superoxide dismutases (SodA and SodC), which neutralize these reactive species and aid in bacterial survival (Shastri et al., 2018).

In addition to ROS, cytokine production plays a crucial role in TB immunity. Cytokines such as interleukin-4 (IL-4) and interleukin-17a (IL-17a) are key mediators in the immune response to *M. tuberculosis*. IL-4, produced by Th2 cells, is involved in the regulation of the immune response and can modulate Th1 responses, which are essential for controlling *M. tuberculosis* infection. While IL-4 may provide protection against excessive inflammation in some contexts, its elevated levels in TB have been associated with enhanced immunopathology and disease progression (Pooran et al., 2019). On the other hand, IL-17a, a cytokine produced by Th17 cells, plays a role in neutrophil recruitment and granuloma formation, which are essential for controlling *M. tuberculosis* growth. However, excessive IL-17a production can lead to immunopathological damage, promoting tissue damage and cavitation (Torrado et al., 2010).

Understanding the regulation of these cytokines in the context of drug-resistant *M. tuberculosis* strains is critical for developing new therapeutic strategies.

The interaction between *M. tuberculosis* and the host immune system is influenced not only by the host’s immune status but also by the strain of *M. tuberculosis* involved. Drug-resistant strains, such as MDR and XDR isolates, may evoke different immune responses compared to drug-sensitive strains (Geffner et al., 2009). Previous studies have shown that drug-resistant *M. tuberculosis* strains exhibit altered immune evasion strategies, which may contribute to their pathogenicity and ability to persist in the host. These strains often exhibit increased virulence, potentially due to genetic mutations that affect their interaction with immune cells and the cytokine profile they induce (Bobba et al., 2023). For instance, MDR strains have been reported to induce higher levels of IL-4, potentially exacerbating the inflammatory response and leading to more severe disease (Ranaivomanana et al., 2021). However, studies investigating the comparative immune responses to drug-resistant and drug-sensitive *M. tuberculosis* strains, particularly in the context of live mycobacterial stimulation, are limited.

Most studies to date have used laboratory-adapted strains of *M. tuberculosis* (H37Rv) or in vitro antigens to study immune responses (Devasundaram et al., 2016; Li et al., 2017; Li et al., 2018). While these studies have provided valuable insights, they do not fully replicate the natural host-pathogen interactions that occur during active infection. The use of live clinical isolates, particularly those with different drug resistance profiles, offers a more accurate reflection of the immune responses encountered during infection. Live bacteria are more likely to trigger authentic immune responses, including cytokine production, cell proliferation, and ROS production, which are critical for controlling the infection and determining the outcome of the disease.

This study aims to evaluate the cellular immune responses of peripheral blood mononuclear cells (PBMCs) from purified protein derivative (PPD)-positive and PPD-negative healthy volunteers upon stimulation with live clinical isolates of *M. tuberculosis*. The study focuses on cell proliferation, cytokine production (IL-4 and IL-17a), and ROS production, with comparisons made between drug-sensitive and drug-resistant isolates, including MDR and streptomycin-independent (SI) resistant strains. By examining these immune parameters in response to live *M. tuberculosis* strains, the study seeks to better understand the immune mechanisms underlying drug-resistant TB and the differences in host responses that may contribute to the pathogenesis of these strains.

## Methodology

### Study Subjects

A tuberculin skin test using 5 TU of tuberculin was conducted on healthy volunteers who had been vaccinated with Bacille Calmette-Guérin (BCG) and had no history of clinical tuberculosis infection. Ten individuals with positive tuberculin skin test results (PPD+) and six with negative results (PPD−) were included in the study. The volunteers were screened to ensure the absence of chronic illnesses, no known contact with TB patients, and no acute medical conditions at the time of enrollment. Heparinized whole blood samples were collected from all participants.

### M. tuberculosis Isolates and Growth Conditions

Clinical *M. tuberculosis* isolates used in the study included multidrug-resistant (MDR) isolates resistant to isoniazid and rifampicin, streptomycin-isoniazid-resistant isolates, drug-sensitive isolates (susceptible to isoniazid, rifampicin, streptomycin, and ethambutol), and the standard strain H37Rv. These isolates were subcultured from Mycobacteria Growth Indicator Tubes (MGIT) onto Löwenstein–Jensen (LJ) medium and incubated at 37°C for 2–3 weeks. A single-cell suspension was prepared by transferring a 10 mg moist-weight cell pellet into a hard glass bottle containing sterile distilled water and glass beads. The suspension was vortexed to emulsify and allowed to settle before adding RPMI media containing 1% antibiotic-antimycotic mixture and 10% fetal bovine serum (FBS). After standing for 15 minutes, the top 9 mL of the suspension was transferred to a fresh tube for subsequent stimulation experiments.

### Peripheral Blood Mononuclear Cell (PBMC) Isolation and Stimulation

PBMCs were isolated from heparinized blood using density gradient centrifugation with Histo-Paque as per established protocols (Birkness et al., 2007). After layering blood diluted with RPMI medium over Histo-Paque, samples were centrifuged at 400×g for 30 minutes at 25°C. The mononuclear cell layer was harvested, washed twice with RPMI, and resuspended in RPMI medium with 10% dimethyl sulfoxide (DMSO) for storage. Prior to stimulation, PBMC viability and live cell counts were assessed using the trypan blue exclusion method (Turner et al., 1996). PBMCs were plated at a density of 1 × 10^6^ cells/well in 24-well plates and stimulated with single-cell suspensions of M. tuberculosis isolates at a multiplicity of infection (MOI) of 5. Phytohemagglutinin (PHA) served as a positive control, while unstimulated PBMCs in RPMI medium served as a negative control. The plates were incubated at 37°C with 5% CO_2_ for 24, 48, 120, and 216 hours. After incubation, the well contents were collected, centrifuged at 4000 rpm for 15 minutes at 10°C, and the pellet and supernatant were stored separately at −80°C until further analysis.

### PBMC Proliferation Assay

Cell proliferation was quantified using a WST-8 cell proliferation assay kit (Cayman Chemical, Ann Arbor, MI, USA). Known cell densities ranging from 1 × 10^6^ to 7 × 10^6^ PBMCs/well in RPMI medium were used to establish a standard curve (Mathur et al., 2014). Stimulated cells were plated alongside standards, and 10 µL of WST-8 reagent was added to each well. Plates were incubated for 4 hours at 37°C in a CO_2_ incubator, and absorbance was measured at 450 nm using a microplate reader. Proliferation rates were determined based on the standard curve and compared across PPD+ and PPD− groups, as well as between resistant and sensitive isolates.

### Cytokine Estimation by ELISA

Quantification of IL-4 and IL-17a cytokines was performed using a commercial ELISA kit (Peprotech ELISA Development Kit). Supernatant from stimulated PBMCs was used for the assay. ELISA plates were coated with capture antibody overnight at 4°C, washed, and blocked before adding standards and test samples. Biotinylated secondary antibodies, avidin-HRP conjugate, and substrate solution were sequentially added with incubation and washing steps between each addition. Absorbance was measured at 405 nm with wavelength correction at 650 nm. Cytokine concentrations were calculated using a four-parameter regression curve.

### Reactive Oxygen Species (ROS) Assay

The production of ROS by PBMCs was measured using the DCFDA method (Shinomol et al., 2008). Cells were stained with 1 mM DCFDA in Locke’s buffer, incubated at 37°C for 10 minutes in the dark, and washed twice with phosphate-buffered saline (PBS). After resuspension in lysis buffer and centrifugation at 13,000 rpm, the supernatant was analyzed for fluorescence using a plate reader with excitation and emission wavelengths of 430 and 530 nm, respectively. ROS activity was expressed as fluorescence intensity per microgram of protein, with protein concentrations determined by Bradford’s assay.

### Statistical Analysis

Data were analyzed using GraphPad online software. Quantitative variables were compared between groups using the Mann–Whitney U test, with statistical significance set at p < 0.05. Results were expressed as means or ranges, and data were presented graphically where applicable.

## Results

### Analysis of PBMC Proliferation

The effect of *M. tuberculosis* clinical isolates on PBMC proliferation was assessed using the WST-8 assay at various time points (Day 1, 2, 5, and 9 post-infection). Proliferative responses among PPD− and PPD+ healthy volunteers are represented in Table 1. Among the isolates, the streptomycin-isoniazid (SI) resistant isolate induced the highest proliferation in both PPD− and PPD+ samples. While proliferation was observed on Day 1, the differences were not statistically significant. On Day 2, a significant increase in proliferation was observed for the SI resistant isolate among PPD− and PPD+ samples (P = 0.0005) and in comparison, with the sensitive isolate for both PPD− (P = 0.0005) and PPD+ (P < 0.0001). By Day 5, the multidrug-resistant (MDR) isolate showed a significant difference between PPD− and PPD+ samples (P = 0.0034), although there was no significant change compared to the sensitive isolate. The SI resistant isolate, however, displayed a significant difference in comparison to the sensitive isolate for both PPD− (P = 0.0013) and PPD+ (P < 0.0001). On Day 9, proliferation responses declined compared to Day 5. Nevertheless, the MDR isolate exhibited a significant difference between PPD− and PPD+ samples (P = 0.0168), while the SI resistant isolate maintained significant differences compared to the sensitive isolate for both PPD− (P = 0.0004) and PPD+ (P < 0.0001).

**Table 1:** Cell proliferation among PPD+ and PPD-healthy volunteers after infection with *M. tuberculosis*. H37Rv: Standard strain; Sen: Sensitive isolate; SI: Streptomycin-isoniazid resistant isolate; MDR: Multidrug resistant isolate; PHA: Phytohaemagglutinin (Positive control); Uns: Unstimulated PBMCs (negative control). The symbol ^*^ represents the statistically significant difference between PPD- and PPD+; # represents a statistically significant difference of resistant isolate with sensitive isolate.

|  | Day 1 |  | Day 2 |  | Day 5 |  | Day 9 |  |
| --- | --- | --- | --- | --- | --- | --- | --- | --- |
| | PPD-<br>Mean ± SD<br>(Range) | PPD+<br>Mean ± SD<br>(Range) | PPD-<br>Mean ± SD<br>(Range) | PPD+<br>Mean ± SD<br>(Range) | PPD-<br>Mean ± SD<br>(Range) | PPD+<br>Mean ± SD<br>(Range) | PPD-<br>Mean ± SD<br>(Range) | PPD+<br>Mean ± SD<br>(Range) |
| **H37Rv** | 1.33 ± 0.87<br>(0.61-2.30) | 1.52 ± 0.54<br>(1.03-2.39) | 1.53 ± 0.37<br>(1.12-1.84) | 3.28 ± 0.91<br>(2.15-4.43) | 2.93 ± 0.62<br>(2.52-3.65) | 4.94 ± 2.21<br>(3.16-8.72) | 2.14 ± 0.45<br>(1.65-2.53) | 4.31 ± 2.63<br>(1.89-8.69) |
| **Sen** | 1.70 ± 0.79<br>(0.95-2.53) | 2.88 ± 0.94<br>(1.87-4.18) | 2.54 ± 0.47<br>(2.21-3.08) | 2.92 ± 2.17<br>(0.82-5.90) | 3.8 ± 2.21<br>(1.25-5.17) | 5.71 ± 4.01<br>(1.65-11.38) | 2.95 ± 1.99<br>(1.40-5.19) | 3.52 ± 1.55<br>(1.20-5.23) |
| **SI** | 5.95 ± 0.29<br>(5.71-6.27) | 5.40 ± 2.19<br>(3.26-8.69) | 7.27 ± 4.20*#<br>(2.53-10.53) | 6.64 ± 0.46*#<br>(6.04-7.29) | 12.92 ± 2.20#<br>(10.51-14.83) | 12.47 ± 1.89#<br>(10.11-14.70) | 7.83 ± 5.33#<br>(3.86-13.89) | 10.2 ± 2.43#<br>(7.85-14.1) |
| **MDR** | 1.49 ± 0.16<br>(1.33-1.64) | 2.76 ± 1.13<br>(1.94-4.76) | 2.90 ± 1.39<br>(1.87-4.48) | 3.35 ± 2.44<br>(1.48-7.39) | 3.65 ± 1.10*<br>(2.72-4.87) | 4.22 ± 1.11*<br>(3.09-5.80) | 3.12 ± 0.62*<br>(2.73-3.84) | 3.80 ± 1.84*<br>(2.73-7.08) |
| **PHA** | 1.05 ± 0.71<br>(0.51-1.86) | 1.39 ± 1.06<br>(0.65-3.24) | 1.11 ± 0.23<br>(0.86-1.30) | 1.48 ± 1.35<br>(0.12-3.14) | 2.15 ± 0.61<br>(1.45-2.59) | 3.00 ± 2.44<br>(1.30-7.21) | 2.32 ± 1.08<br>(1.53-3.55) | 2.16 ± 1.33<br>(0.85-4.19) |
| **UnS** | 0.8 ± 0.15<br>(0.70-0.97) | 0.96 ± 0.79<br>(0.31-2.32) | 1.75 ± 0.72<br>(1.07-2.50) | 1.28 ± 0.55<br>(0.72-2.08) | 1.83 ± 0.37<br>(1.45-2.18) | 2.02 ± 1.53<br>(1.05-4.65) | 1.76 ± 0.18<br>(1.60-1.95) | 1.83 ± 0.93<br>(0.92-3.25) |

### IL-4 Cytokine Analysis

The levels of IL-4 cytokine in PBMC supernatants at Days 1, 2, 5, and 9 post-infection are shown in Figure 1A and 1B. On Day 1, IL-4 concentrations were comparable across all isolates. By Day 2, the SI resistant isolate exhibited significant differences between PPD− and PPD+ samples (P = 0.0034), as did the MDR isolate (P = 0.0013). Compared to the sensitive isolate, significant differences were observed for the SI resistant isolate in PPD− samples (P = 0.0004) and the MDR isolate in both PPD− (P = 0.0167) and PPD+ (P = 0.0159). On Day 5, significant differences were noted between the SI resistant and sensitive isolates (P = 0.0403) and between the MDR and sensitive isolates (P = 0.0403) in PPD+ samples. On Day 9, the MDR isolate demonstrated significant differences between PPD− and PPD+ samples (P = 0.0034) and compared to the sensitive isolate in both PPD− (P = 0.0004) and PPD+ (P = 0.0297).

**Figure 1:**
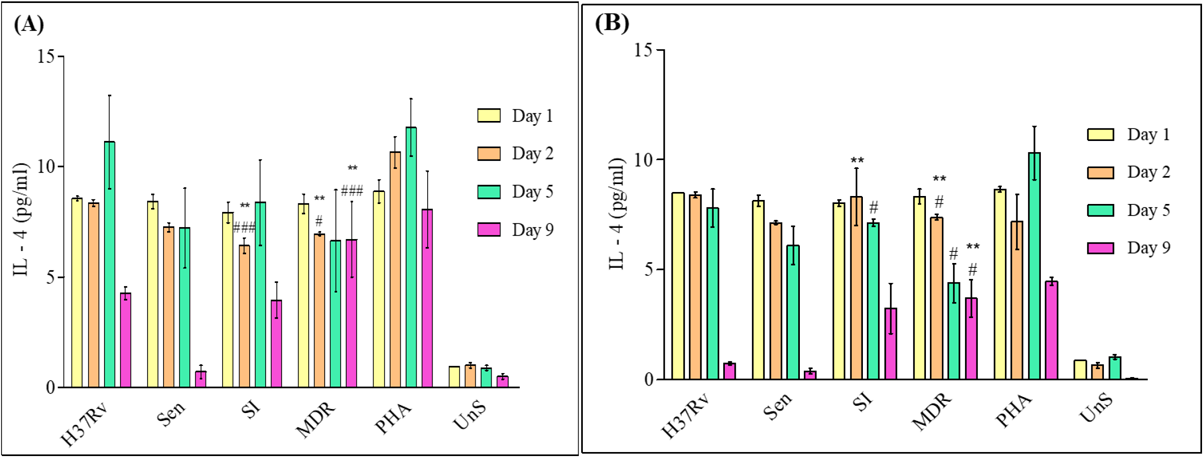
IL-4 cytokine concentration among PPD- (A) and PPD+ (B) healthy volunteers after infection with *M. tuberculosis*. The y-axis represents IL-4 concentration (Mean ± SD) values. The x-axis represents different M. tuberculosis isolates used for infection. The symbol ^*^ represents the statistically significant difference between PPD- and PPD+; # represents a statistically significant difference of resistant isolate with sensitive isolate. H37Rv: Standard strain; Sen: Sensitive isolate; SI: Streptomycin-isoniazid resistant isolate; MDR: Multidrug resistant isolate; PHA: Phytohaemagglutinin (Positive control); Uns: Unstimulated PBMCs (negative control).

### IL-17a Cytokine Analysis

IL-17a levels measured in PBMC supernatants at Days 1, 2, 5, and 9 are presented in Figures 2A and 2B. On Day 1, IL-17a concentrations showed no significant differences across isolates. By Day 2, significant differences were observed between PPD− and PPD+ samples for the sensitive isolate (P = 0.0034) and for the MDR isolate compared to the sensitive isolate in PPD+ samples (P < 0.0001). On Day 5, significant differences in IL-17a levels were seen for the sensitive isolate (P = 0.032), the SI resistant isolate (P = 0.0168), and the MDR isolate (P = 0.0074) between PPD− and PPD+ samples. However, no significant differences were noted between resistant isolates and the sensitive isolate in both PPD− and PPD+ groups. On Day 9, the sensitive isolate showed significant differences between PPD− and PPD+ samples (P = 0.017), while resistant isolates displayed no significant changes compared to the sensitive isolate.

**Figure 2:**
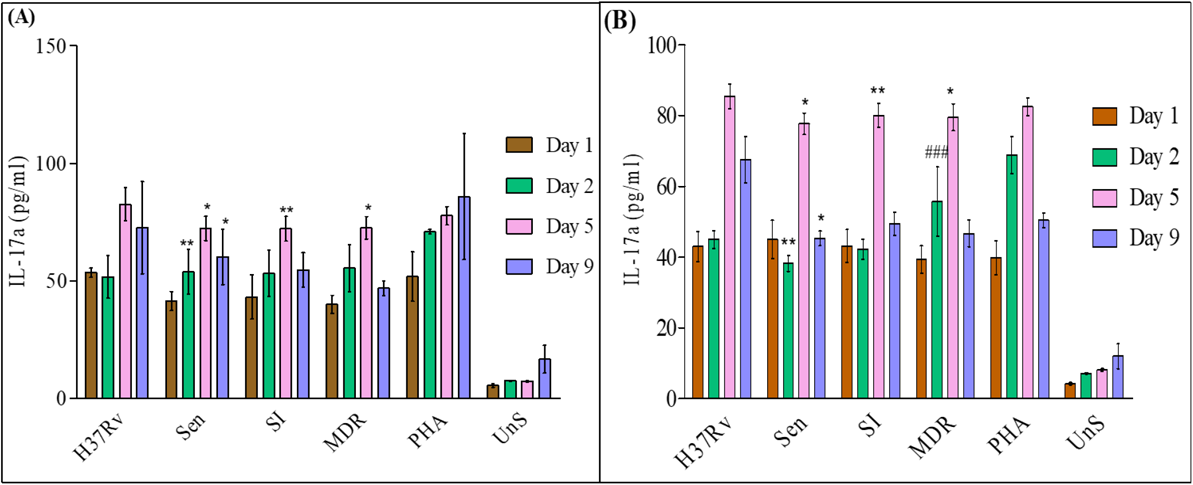
IL-17a cytokine concentration among PPD- (A) and PPD+ (B) healthy volunteers after infection with *M. tuberculosis*. The y-axis represents IL-17a concentration (Mean ± SD) values. The x-axis represents different M. tuberculosis isolates used for infection. The symbol ^*^ represents the statistically significant difference between PPD- and PPD+; # represents a statistically significant difference of resistant isolate with sensitive isolate. H37Rv: Standard strain; Sen: Sensitive isolate; SI: Streptomycin-isoniazid resistant isolate; MDR: Multidrug resistant isolate; PHA: Phytohaemagglutinin (Positive control); Uns: Unstimulated PBMCs (negative control).

### ROS Production Analysis

Reactive oxygen species (ROS) production in PBMCs after infection with *M. tuberculosis* isolates was assessed using the DCFDA method. The response to infection varied among clinical isolates, as shown in Figures 3A and 3B. No significant differences were observed between PPD− and PPD+ samples across all isolates until the last day of incubation. On Day 1, ROS production was significantly elevated for the SI resistant isolate compared to the sensitive isolate in PPD+ samples (P = 0.0054). On Day 2, the MDR isolate showed significant differences compared to the sensitive isolate in PPD+ samples (P < 0.0001). By Day 5, the SI resistant isolate displayed significant differences compared to the sensitive isolate in both PPD− (P = 0.0059) and PPD+ (P = 0.0054), and the MDR isolate showed significant differences in PPD+ samples (P = 0.0417). On Day 9, no significant differences were observed among clinical isolates.

**Figure 3:**
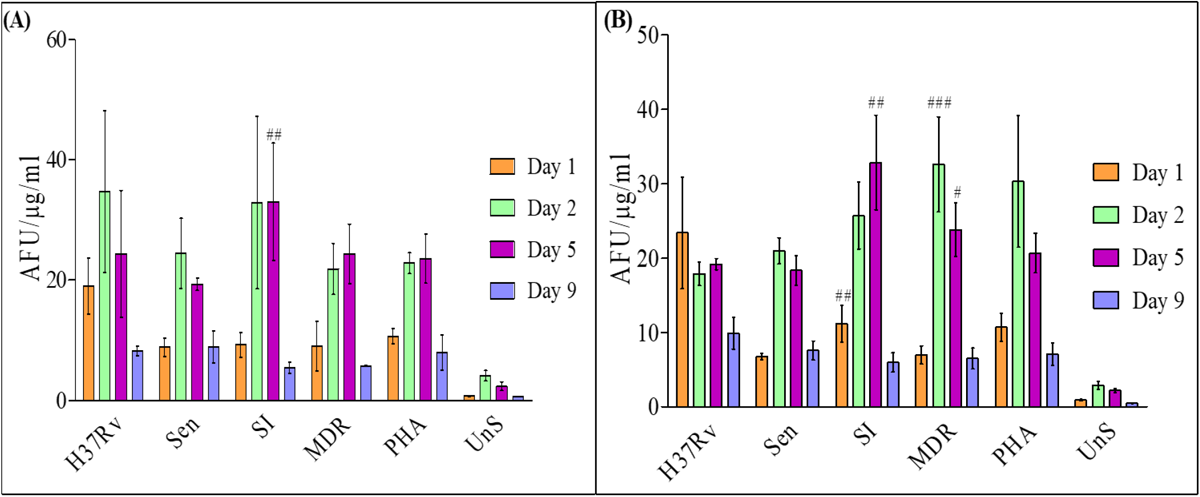
ROS production among PPD- (A) and PPD+ (B) healthy volunteers after infection with *M. tuberculosis*. The graph represents Mean ± SD values. Y-axis represents average fluorescent unit (AFU)/ µg of protein/ ml. The x-axis represents different M. tuberculosis isolates used for infection. H37Rv: Standard strain; Sen: Sensitive isolate; SI: Streptomycin-isoniazid resistant isolate; MDR: Multidrug resistant isolate; PHA: Phytohaemagglutinin (Positive control); Uns: Unstimulated PBMCs (negative control). The symbol # represents a statistically significant difference of resistant isolate with sensitive isolate.

## Discussion

### Assessment of Immune Response

The study evaluated cellular immune responses using PBMCs from PPD- and PPD+ healthy volunteers exposed to live *M. tuberculosis* clinical isolates, including MDR, SI-resistant, sensitive isolates, and the standard H37Rv strain. Live organisms were used as they closely mimic in vivo host infection, where immune cells encounter intact bacteria. This approach provides insights into immune modulation in response to different isolates, particularly given the observed changes in cell wall morphology. Understanding immune responses to resistant isolates offers critical information on protective and pathological mechanisms in TB, aiding in developing effective therapies and vaccines.

### PBMC Proliferation

Cell proliferation, an indicator of cell-mediated immunity, was higher in PPD+ than PPD- individuals after exposure to live *M. tuberculosis*, likely due to sensitized memory T cells in PPD+ subjects (Shahemabadi et al., 2007). Interestingly, the SI-resistant isolate induced a stronger proliferative response in PPD- individuals compared to PPD+, potentially due to cross-reactive epitopes or impaired immune recognition in PPD+ subjects. These findings align with prior studies demonstrating similar proliferative responses between MDR and sensitive isolates in PPD+ subjects (Basile et al., 2011). The observed decline in proliferation by day 9 likely reflects cytolytic activity or immune exhaustion (Turner et al., 1996). The unique response to SI-resistant isolates warrants further investigation to identify specific epitopes contributing to this phenomenon.

### IL-4 Secretion

IL-4, a cytokine implicated in TB progression and immune modulation, showed low overall levels in this study, consistent with reports attributing this to the cytokine’s short half-life or low mRNA copy number (Rook et al., 2004). In PPD+ individuals, IL-4 levels increased by day 2 but declined by day 9, suggesting a protective shift away from Th2 responses towards a Th1-dominant immune profile. Conversely, resistant isolates, particularly MDR strains, induced higher IL-4 levels than sensitive isolates. Elevated IL-4 production may contribute to immune deviation, impairing protective Th1 responses and promoting T cell apoptosis, which is linked to increased disease severity (Minty et al., 1997; Shekar-Abi et al., 2004). These findings reinforce the pathogenic role of MDR strains through immune dysregulation.

### IL-17a Secretion

IL-17a, a cytokine critical for granuloma formation and neutrophil-mediated inflammation, increased significantly in all samples by day 5, followed by a decline on day 9. PPD+ individuals exhibited stronger IL-17a responses, reflecting their sensitized immune state and effective memory T cell activation (Umemura et al., 2007). MDR isolates elicited significantly higher IL-17a levels compared to sensitive isolates, suggesting a heightened inflammatory response potentially associated with tissue damage (Cruz et al., 2006). While IL-17a contributes to protective immunity, excessive production may exacerbate immunopathology. Balancing Th1 and Th17 responses is critical to controlling bacterial growth while minimizing host tissue damage (Basile et al., 2011).

### ROS Production

ROS production, a component of the innate immune response, showed no significant differences between PPD- and PPD+ individuals, underscoring its nonspecific nature. However, resistant isolates, particularly MDR strains, induced significantly higher ROS levels compared to sensitive isolates. This may be attributed to impaired ROS-neutralizing mechanisms, such as mutations in genes encoding ROS-scavenging enzymes (e.g., katG, ahpC) (Ng et al., 2004; Master et al. 2002). Prior studies have linked such mutations to oxidative stress susceptibility, supporting the hypothesis that resistance mechanisms compromise the pathogen’s ability to counteract host oxidative defenses (Kohanski et al., 2010). Targeting these vulnerabilities may provide novel therapeutic strategies for resistant TB strains.

### Clinical Implications

This study highlights distinct immune responses elicited by resistant and sensitive *M. tuberculosis* isolates. Resistant isolates, particularly MDR strains, are associated with heightened IL-4 and IL-17a production and impaired ROS detoxification, contributing to immune dysregulation and severe pathogenesis. These findings underscore the importance of understanding host-pathogen interactions to inform vaccine design and immunotherapeutic approaches. Future studies should explore in vivo models to validate these immune dynamics and assess potential interventions targeting immune deviations caused by resistant strains.

### Limitations and Future Directions

The study’s limitations include the use of in vitro PBMC cultures, which may not fully replicate in vivo immune responses. Additionally, the short-term nature of the study precludes analysis of long-term immune dynamics. Future research should incorporate animal models or human infection studies to validate these findings and explore multi-cytokine profiling to provide a more comprehensive view of immune modulation.

## Conclusion

The pathogenesis of tuberculosis (TB) involves complex interactions between the host immune system, the pathogen, and environmental factors. This study uniquely evaluates cell proliferation, cytokine induction, and reactive oxygen species (ROS) production in host cells stimulated with live *M. tuberculosis* isolates, including drug-resistant strains. Using clinical isolates provides realistic insights into immune responses, highlighting significant variations between sensitive and resistant strains. While the study underscores genotype-environment interactions in drug resistance, it remains unclear if these changes are drug-induced. These findings emphasize the need for further research to understand the immune modulation in drug-resistant TB and its clinical implications.

## References

1. Alsayed SSR, Gunosewoyo H. Tuberculosis: Pathogenesis, Current Treatment Regimens and New Drug Targets. Int J Mol Sci. 2023 Mar 8;24(6):5202. doi: 10.3390/ijms24065202. PMID: 36982277; PMCID: PMC10049048.

2. Bagcchi S. WHO’s Global Tuberculosis Report 2022. Lancet Microbe. 2023 Jan;4(1):e20. doi: 10.1016/S2666-5247(22)00359-7. Epub 2022 Dec 12. PMID: 36521512.

3. Lange C, Abubakar I, Alffenaar JW, Bothamley G, Caminero JA, Carvalho AC, Chang KC, Codecasa L, Correia A, Crudu V, Davies P, Dedicoat M, Drobniewski F, Duarte R, Ehlers C, Erkens C, Goletti D, Günther G, Ibraim E, Kampmann B, Kuksa L, de Lange W, van Leth F, van Lunzen J, Matteelli A, Menzies D, Monedero I, Richter E, Rüsch-Gerdes S, Sandgren A, Scardigli A, Skrahina A, Tortoli E, Volchenkov G, Wagner D, van der Werf MJ, Williams B, Yew WW, Zellweger JP, Cirillo DM; TBNET. Management of patients with multidrug-resistant/extensively drug-resistant tuberculosis in Europe: a TBNET consensus statement. Eur Respir J. 2014 Jul;44(1):23–63. doi: 10.1183/09031936.00188313. Epub 2014 Mar 23. PMID: 24659544; PMCID: PMC4076529.

4. Sachan RSK, Mistry V, Dholaria M, Rana A, Devgon I, Ali I, Iqbal J, Eldin SM, Mohammad Said Al-Tawaha AR, Bawazeer S, Dutta J, Karnwal A. Overcoming Mycobacterium tuberculosis Drug Resistance: Novel Medications and Repositioning Strategies. ACS Omega. 2023 Sep 1;8(36):32244–32257. doi: 10.1021/acsomega.3c02563. PMID: 37720746; PMCID: PMC10500578.

5. de Martino M, Lodi L, Galli L, Chiappini E. Immune Response to Mycobacterium tuberculosis: A Narrative Review. Front Pediatr. 2019 Aug 27;7:350. doi: 10.3389/fped.2019.00350. PMID: 31508399; PMCID: PMC6718705.

6. Lerner TR, Borel S, Gutierrez MG. The innate immune response in human tuberculosis. Cell Microbiol. 2015 Sep;17(9):1277–85. doi: 10.1111/cmi.12480. Epub 2015 Jul 28. PMID: 26135005; PMCID: PMC4832344.

7. Juan-García J, García-García S, Guerra-Laso JM, Raposo-García S, Diez-Tascón C, Nebreda-Mayoral T, López-Fidalgo E, López-Medrano R, Fernández-Maraña A, Rivero-Lezcano OM. In vitro infection with Mycobacterium tuberculosis induces a distinct immunological pattern in blood from healthy relatives of tuberculosis patients. Pathog Dis. 2017 Nov 30;75(8). doi: 10.1093/femspd/ftx109. PMID: 29048475.

8. Shastri MD, Shukla SD, Chong WC, Dua K, Peterson GM, Patel RP, Hansbro PM, Eri R, O’Toole RF. Role of Oxidative Stress in the Pathology and Management of Human Tuberculosis. Oxid Med Cell Longev. 2018 Oct 11;2018:7695364. doi: 10.1155/2018/7695364. PMID: 30405878; PMCID: PMC6201333.

9. Pooran A, Davids M, Nel A, Shoko A, Blackburn J, Dheda K. IL-4 subverts mycobacterial containment in Mycobacterium tuberculosis-infected human macrophages. Eur Respir J. 2019 Aug 8;54(2):1802242. doi: 10.1183/13993003.02242-2018. PMID: 31097521.

10. Torrado E, Cooper AM. IL-17 and Th17 cells in tuberculosis. Cytokine Growth Factor Rev. 2010 Dec;21(6):455–62. doi: 10.1016/j.cytogfr.2010.10.004. Epub 2010 Nov 12. PMID: 21075039; PMCID: PMC3032416.

11. Geffner L, Yokobori N, Basile J, Schierloh P, Balboa L, Romero MM, Ritacco V, Vescovo M, González Montaner P, Lopez B, Barrera L, Alemán M, Abatte E, Sasiain MC, de la Barrera S. Patients with multidrug-resistant tuberculosis display impaired Th1 responses and enhanced regulatory T-cell levels in response to an outbreak of multidrug-resistant Mycobacterium tuberculosis M and Ra strains. Infect Immun. 2009 Nov;77(11):5025–34. doi: 10.1128/IAI.00224-09. Epub 2009 Aug 31. PMID: 19720756; PMCID: PMC2772532.

12. Bobba S, Khader SA. Rifampicin drug resistance and host immunity in tuberculosis: more than meets the eye. Trends Immunol. 2023 Sep;44(9):712–723. doi: 10.1016/j.it.2023.07.003. Epub 2023 Aug 3. PMID: 37543504; PMCID: PMC11170062.

13. Ranaivomanana P, Rabodoarivelo MS, Ndiaye MDB, Rakotosamimanana N, Rasolofo V. Different PPD-stimulated cytokine responses from patients infected with genetically distinct Mycobacterium tuberculosis complex lineages. Int J Infect Dis. 2021 Mar;104:725–731. doi: 10.1016/j.ijid.2021.01.073. Epub 2021 Feb 5. PMID: 33556615.

14. Devasundaram S, Gopalan A, Das SD, Raja A. Proteomics Analysis of Three Different Strains of Mycobacterium tuberculosis under In vitro Hypoxia and Evaluation of Hypoxia Associated Antigen’s Specific Memory T Cells in Healthy Household Contacts. Front Microbiol. 2016 Sep 9;7:1275. doi: 10.3389/fmicb.2016.01275. PMID: 27667981; PMCID: PMC5017210.

15. Li H, Wang XX, Wang B, Fu L, Liu G, Lu Y, Cao M, Huang H, Javid B. Latently and uninfected healthcare workers exposed to TB make protective antibodies against Mycobacterium tuberculosis. Proc Natl Acad Sci U S A. 2017 May 9;114(19):5023–5028. doi: 10.1073/pnas.1611776114. Epub 2017 Apr 24. PMID: 28438994; PMCID: PMC5441709.

16. Li H, Javid B. Antibodies and tuberculosis: finally coming of age? Nat Rev Immunol. 2018 Sep;18(9):591–596. doi: 10.1038/s41577-018-0028-0. PMID: 29872140.

17. Birkness KA, Guarner J, Sable SB, Tripp RA, Kellar KL, Bartlett J, Quinn FD. An in vitro model of the leukocyte interactions associated with granuloma formation in Mycobacterium tuberculosis infection. Immunol Cell Biol. 2007 Feb-Mar;85(2):160–8. doi: 10.1038/sj.icb.7100019. Epub 2007 Jan 2. PMID: 17199112.

18. Turner J, Dockrell HM. Stimulation of human peripheral blood mononuclear cells with live Mycobacterium bovis BCG activates cytolytic CD8+ T cells in vitro. Immunology. 1996 Mar;87(3):339–42. doi: 10.1046/j.1365-2567.1996.512590.x. PMID: 8778016; PMCID: PMC1384099.

19. Mathur A, Abd Elmageed ZY, Liu X, Kostochka ML, Zhang H, Abdel-Mageed AB, Mondal D. Subverting ER-stress towards apoptosis by nelfinavir and curcumin coexposure augments docetaxel efficacy in castration resistant prostate cancer cells. PLoS One. 2014 Aug 14;9(8):e103109. doi: 10.1371/journal.pone.0103109. PMID: 25121735; PMCID: PMC4133210.

20. Shinomol GK Muralidhara. Prophylactic neuroprotective property of Centella asiatica against 3-nitropropionic acid induced oxidative stress and mitochondrial dysfunctions in brain regions of prepubertal mice. Neurotoxicology. 2008 Nov;29(6):948–57. doi: 10.1016/j.neuro.2008.09.009. Epub 2008 Sep 27. PMID: 18930762.

21. Shahemabadi AS, Hosseini AZ, Shaghsempour S, Masjedi MR, Rayani M, Pouramiri M. Evaluation of T cell immune responses in multi-drug-resistant tuberculosis (MDR-TB) patients to Mycobacterium tuberculosis total lipid antigens. Clin Exp Immunol. 2007 Aug;149(2):285–94. doi: 10.1111/j.1365-2249.2007.03406.x. Epub 2007 May 9. PMID: 17490401; PMCID: PMC1941963.

22. Basile JI, Geffner LJ, Romero MM, Balboa L, Sabio Y García C, Ritacco V, García A, Cuffré M, Abbate E, López B, Barrera L, Ambroggi M, Alemán M, Sasiain MC, de la Barrera SS. Outbreaks of mycobacterium tuberculosis MDR strains induce high IL-17 T-cell response in patients with MDR tuberculosis that is closely associated with high antigen load. J Infect Dis. 2011 Oct 1;204(7):1054–64. doi: 10.1093/infdis/jir460. PMID: 21881121.

23. Rook GA, Hernandez-Pando R, Dheda K, Teng Seah G. IL-4 in tuberculosis: implications for vaccine design. Trends Immunol. 2004 Sep;25(9):483–8. doi: 10.1016/j.it.2004.06.005. PMID: 15324741.

24. Minty A, Asselin S, Bensussan A, Shire D, Vita N, Vyakarnam A, Wijdenes J, Ferrara P, Caput D. The related cytokines interleukin-13 and interleukin-4 are distinguished by differential production and differential effects on T lymphocytes. Eur Cytokine Netw. 1997 Jun;8(2):203–13. PMID: 9262969.

25. Shekar-Abi M, Miandehi N, Mansoori SD, Tavasoli Fayri M, Alibahar M, Amirkhani A, Mirsaeidi SM. The study of Th1/Th2 cytokine profiles (IL-10, IL-12, IL-4, and IFNγ) in PBMCs of patients with multidrug resistant tuberculosis and newly diagnosed drug responsive cases. TANAFFOS (Respiration). 2004 Jun 1;3(2 (spring)):25–31.

26. Umemura M, Yahagi A, Hamada S, Begum MD, Watanabe H, Kawakami K, Suda T, Sudo K, Nakae S, Iwakura Y, Matsuzaki G. IL-17-mediated regulation of innate and acquired immune response against pulmonary Mycobacterium bovis bacille Calmette-Guerin infection. J Immunol. 2007 Mar 15;178(6):3786–96. doi: 10.4049/jimmunol.178.6.3786. PMID: 17339477.

27. Cruz A, Khader SA, Torrado E, Fraga A, Pearl JE, Pedrosa J, Cooper AM, Castro AG. Cutting edge: IFN-gamma regulates the induction and expansion of IL-17-producing CD4 T cells during mycobacterial infection. J Immunol. 2006 Aug 1;177(3):1416–20. doi: 10.4049/jimmunol.177.3.1416. PMID: 16849446.

28. Ng VH, Cox JS, Sousa AO, MacMicking JD, McKinney JD. Role of KatG catalase-peroxidase in mycobacterial pathogenesis: countering the phagocyte oxidative burst. Mol Microbiol. 2004 Jun;52(5):1291–302. doi: 10.1111/j.1365-2958.2004.04078.x. PMID: 15165233.

29. Master SS, Springer B, Sander P, Boettger EC, Deretic V, Timmins GS. Oxidative stress response genes in Mycobacterium tuberculosis: role of ahpC in resistance to peroxynitrite and stage-specific survival in macrophages. Microbiology (Reading). 2002 Oct;148(Pt 10):3139–3144. doi: 10.1099/00221287-148-10-3139. PMID: 12368447.

30. Kohanski MA, DePristo MA, Collins JJ. Sublethal antibiotic treatment leads to multidrug resistance via radical-induced mutagenesis. Mol Cell. 2010 Feb 12;37(3):311–20. doi: 10.1016/j.molcel.2010.01.003. PMID: 20159551; PMCID: PMC2840266.

